# *amR*: an R package suite to predict antimicrobial resistance in bacterial pathogens

**DOI:** 10.64898/2026.07.10.734579

**Authors:** Abhirupa Ghosh, Evan P Brenner, Emily A Boyer, Alexander P McKim, Charmie K Vang, Ethan P Wolfe, David Mayer, Raymond L Lesiyon, Janani Ravi

## Abstract

**Motivation:** Identifying bacterial antimicrobial resistance (AMR) is critical for diagnostics and treatment, but resistance is a complex trait arising from myriad mechanisms spanning multiple molecular scales. Existing computational approaches often function as black boxes and rarely explore cross-species or multi-drug patterns. We developed *amR*, an integrated R package suite that provides a complete framework from bacterial genome data curation to interpretable AMR predictions, enabling identification of resistance mechanisms across species and drugs.

**Results:** The *amR* R package suite contains three modular packages. *amRdata* downloads genomes and paired antimicrobial susceptibility testing data from BV-BRC and processes them, constructs pangenomes, and extracts features at gene/protein cluster, protein domain, annotated Clusters of Orthologous Groups and ResFinder AMR-associated features, and structural variant scales; data are stored in memory-efficient formats (Parquet, DuckDB). *amRml* trains interpretable machine learning models per species-drug combination, calculates feature importance and performance metrics, and provides rich ground for hypothesis generation and mechanism discovery. *amRviz* provides an interactive Shiny dashboard to explore metadata distributions and model performance across species and drugs, visualize top predictive AMR features, and analyze cross-model patterns across geographic/temporal strata. We apply the suite to *Shigella sonnei*, achieving a median Matthews Correlation Coefficient of 0.89 across 23 drugs and drug classes. With thousands of genomes, multi-scale features, and interpretable models, *amR* provides an accessible, comprehensive framework for AMR research. The *amR* package suite is installable via GitHub (https://github.com/JRaviLab/amR; BSD-3-Clause license).

## Background and Rationale

Despite intensive study, antimicrobial resistance (AMR) remains a worsening health crisis. Understanding the molecular mechanisms that cause bacterial AMR is critical for effective interventions, and multiple computational methods have been developed to accelerate AMR diagnostics and discovery. While some forms of AMR are well-understood and readily identified by molecular diagnostics (e.g., beta-lactamases), novel mechanisms are constantly evolving.^1^ Prior *in silico* research has leveraged sequence similarity or machine learning (ML) with AMR-labeled bacterial genome sequence data to predict AMR.^2–12^ However, this problem remains challenging: complex combinations of core and accessory genes, rare variants, and context-dependent interactions can all contribute uniquely to AMR across different drugs and species. As bacteria evolve and exchange AMR mechanisms with global morbidity and mortality impacts, there is an increasing need for computational solutions that can be applied quickly to new contexts and provide comprehensive insights into AMR through predictive modeling, mechanism identification, and simplified exploration of results. Furthermore, few studies explore AMR prediction across drug classes, species, or multidrug resistance (MDR).

To predict AMR phenotypes and mechanisms, we leverage abundant whole-genome sequence (WGS) and AMR phenotype data across bacterial isolates to train supervised ML models spanning multiple feature scales across drugs and drug classes. Here, we present *amR* (JRaviLab/amR), an R package suite that introduces a multiscale approach in an easy-to-use programmatic interface for all relevant functions to download and process BV-BRC data and metadata, train ML models, and extract the top features for further research, benchmarking, and experimental follow-up.

Existing computational AMR tools broadly fall into two camps: database-anchored approaches (CARD,^13^ ResFinder,^14^ AMRFinderPlus^15^) that are effective but limited to previously characterized resistance mechanisms, and pangenome-based ML methods (pangenomix^7^; PanKA^11^) that extend beyond known antimicrobial resistance genes (ARGs) by finding associations within WGS-scale data. *amR* extends both paradigms through four advances. First, *amR* integrates features across multiple complementary molecular scales – pangenome gene clusters,^16^ protein clusters,^17^ Pfam protein domains,^18^ Clusters of Orthologous Groups^19,20^ (COGs) or ResFinder^14^ feature annotation, and gene-neighborhood structural variants.^16^ This enables systematic interrogation of resistance signatures that may be lost to noise in single-scale approaches. Second, protein domain and COG features provide a biologically principled path toward cross-species AMR inference: unlike individual genes or SNPs, these features are more widely conserved beyond species boundaries, making domain- and COG-level models more species-agnostic and therefore transferable to pathogens without characterized ARGs, a challenge that existing interpretable approaches have not addressed. Third, *amR* pairs this feature architecture with rigorous, multi-layered model evaluation with label-shuffled baselines, geographic and temporal holdouts, cross-drug and cross-class testing, and multiclass MDR models. Fourth, we provide users with a fully interactive visualization experience with a Shiny-based^21^ data dashboard for their specific datasets. This is all available through an accessible R workflow that enables the intentional evaluation of potential mechanisms rather than reporting predictive performance alone. These design choices position *amR* as a one-stop comprehensive framework for both reliable AMR prediction and hypothesis generation in newly sequenced pathogens.

## Implementation

*amR* is an R package suite for processing bacterial genomics datasets with AMR metadata sourced from BV-BRC,^12,22^ training ML models, identifying top AMR genomic predictors, and analyzing these findings via an interactive dashboard (**Figure 1**). *amR* creates bacterial pangenomes per user-specified taxa as the basis for feature extraction. These pangenomes are then abstracted as feature matrices at five molecular scales – pangenome gene clusters, pangenome graph structural variants, protein clusters, COGs (Clusters of Orthologous Groups), and Pfam domains, along with known antimicrobial resistance genes (ARGs) – and encoded as either binarized feature presence/absence or feature counts. Genomes are paired with their corresponding AMR resistance/susceptibility label from lab-validated phenotypes. These labeled matrices serve as inputs to our ML models. *amR* trains a resistant/susceptible classifier model to predict AMR phenotypes per bug, per drug (antibiotic, drug class), and across bugs and drugs. The *amR* package includes well-documented functions and vignettes to reproduce all aforementioned steps, plus additional functions to summarize and visualize data availability by metadata (e.g., MDR status, temporal and geographic information), model performance, and multiscale genomic predictors (genes, proteins, domains, COGs, ARGs, structural variants) of AMR. *amR* is a modular three-package ecosystem for AMR prediction comprised of: (1) *amRdata* (JRaviLab/amRdata) for data curation and feature extraction, (2) *amRml* (JRaviLab/amRml for interpretable ML model training, and (3) *amRviz* (JRaviLab/amRviz) for interactive visualization and exploration of results. This integrated framework provides end-to-end functionality from annotated genomes to AMR predictions.

**Figure 1.**
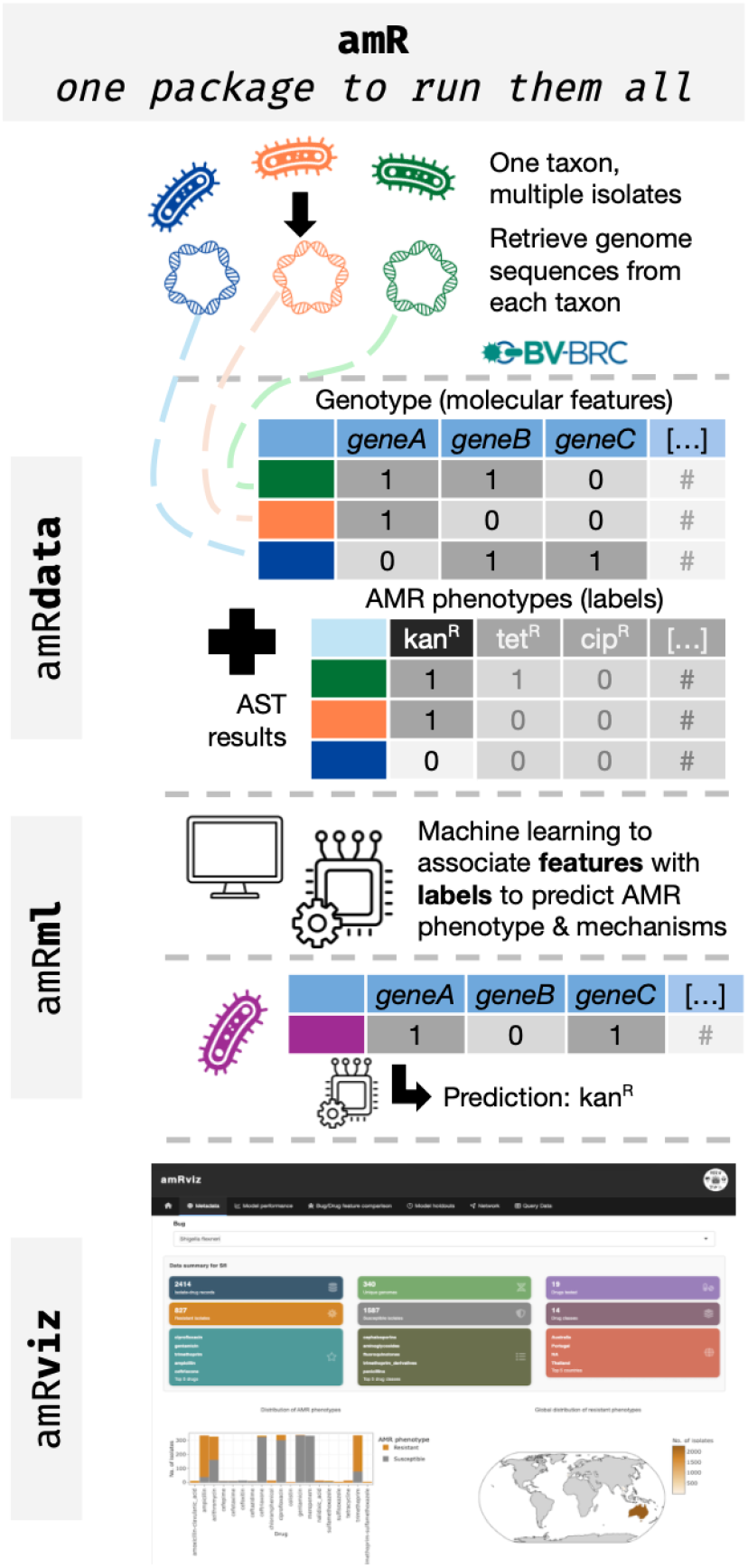
**Overview schematic of the amR package suite** functionality with component packages, *amRdata, amRml*, and *amRviz*.

The *amRdata* package constructs multiscale genomic feature matrices from annotated bacterial genomes with paired antimicrobial susceptibility data to enable AMR prediction. It interfaces with BV-BRC (Bacterial and Viral Bioinformatics Resource Center)^12,22^ through the BV-BRC CLI. Functions query isolate metadata, download genome sequences, retrieve AST results, and apply quality filters. After downloading genomes, *amRdata* extracts features at six scales. Panaroo^16^ constructs graph-based pangenomes, generating gene cluster matrices and identifying structural variants (unique triplets of neighboring genes on the pangenome graph that indicate a genomic rearrangement). For well-sampled species with thousands of isolates, pangenome construction can be parallelized across subsets with results merged. This improves scalability and computational efficiency while enabling analyses that would otherwise be impractical due to memory and runtime constraints. CD-HIT^17^ clusters protein sequences to generate protein cluster-level features. HMMER^23^ queries representative sequences against Hidden Markov Model-based databases for Pfam^24^ domains through InterProScan,^25^ COGs,^19,20^ and ARGs (ResFinder^14^). All matrices and metadata are stored in Parquet format with DuckDB^26^ indexing for efficient querying. External software dependencies use Docker containers for reproducibility and simplified setup.^27^ Finally, *amRdata* enables the summarization and visualization of genomic data and metadata, and supports the exploration of data availability towards training ML models.

The *amRml* package leverages the tidymodels^28^ framework to train interpretable models for AMR prediction. For each drug (and drug class), the package generates input matrices by subsetting isolates with AST data, and creating training/testing splits or cross-validation folds. Logistic regression was chosen as the underlying algorithm for interpretability; coefficients directly indicate feature importance and direction of effect. Models are trained separately for each molecular scale (genes, proteins, domains, COGs, structural variants) using both feature counts and binarized values. After training, *amRml* extracts top features across molecular scales (e.g., *tetA* for tetracycline resistance in *Shigella sonnei*) and calculates performance metrics including balanced accuracy, F1 score, and Matthews correlation coefficient (MCC). The package supports parallel processing across multiple drugs and species. *amRml* includes functionality to evaluate performance and feature robustness using shuffled label matrices and baseline evaluation using Fisher’s exact testing. *amRml* also allows training and cross-testing on stratified and holdout datasets. Additionally, *amRml* implements a multiclass model for predicting resistance to more than one drug class, permitting exploration of the features that could contribute to MDR.

The *amRviz* dashboard is organized into six analysis modules: (i) metadata exploration (geographic distribution, temporal trends, host and isolation source); (ii) model performance comparison across species, drugs, and molecular scales; (iii) feature comparison across bugs and drugs with COG enrichment; (iv) model holdouts, comparing cross-model generalization across geographic and temporal strata; (v) network visualization of drug-feature relationships; and (vi) custom data queries with filtering. Built with Shiny^21^ in a modular architecture, the dashboard renders interactive plots and tables and publication-quality heatmaps, all exportable as CSV or PDF. *amRviz* reads pre-computed Parquet outputs directly, requiring no database backend, and operates directly with *amRdata* and *amRml* outputs.

### Benchmarking

We compared several alternative methods for AMR prediction. The first, PanKA, shares the most similarity with our approach by starting with pangenome-based features for AMR prediction. They demonstrate their performance in selected *E. coli* and *K. pneumoniae* genomes – we used their *K. pneumoniae* genome set as a baseline to compare across methods. We extracted BV-BRC’s AdaBoost *k*-mer models’ predictions for these same genomes. Finally, we used these genomes as input for Kover’s set-covering machine *k*-mer models.

We also note that we were unable to reproduce PanKA’s model runs, so as a workaround, we used their reported performance metric (F1 score) for their specified benchmark set rather than running the models ourselves. In the PanKA benchmark set, some were no longer available through BV-BRC. We used this reduced subset for further model testing. For Kover, phenotype labels and genome data paths were manually formatted for input. BV-BRC’s ML workflow is not designed to be run by users, so we calculated performance on the genomes in our set that had both laboratory-validated and BV-BRC computationally predicted labels.

We therefore compared the performance of BV-BRC, Kover, and PanKA with our amR package models across molecular scales for these ∼1700 *K. pneumoniae* genomes ranging across 11 antibiotics. Our models were the top-performing 9 of 11 times, with binary gene models performing best in 4 of 9 models (**Figure 2; Table S1**).

**Figure 2.**
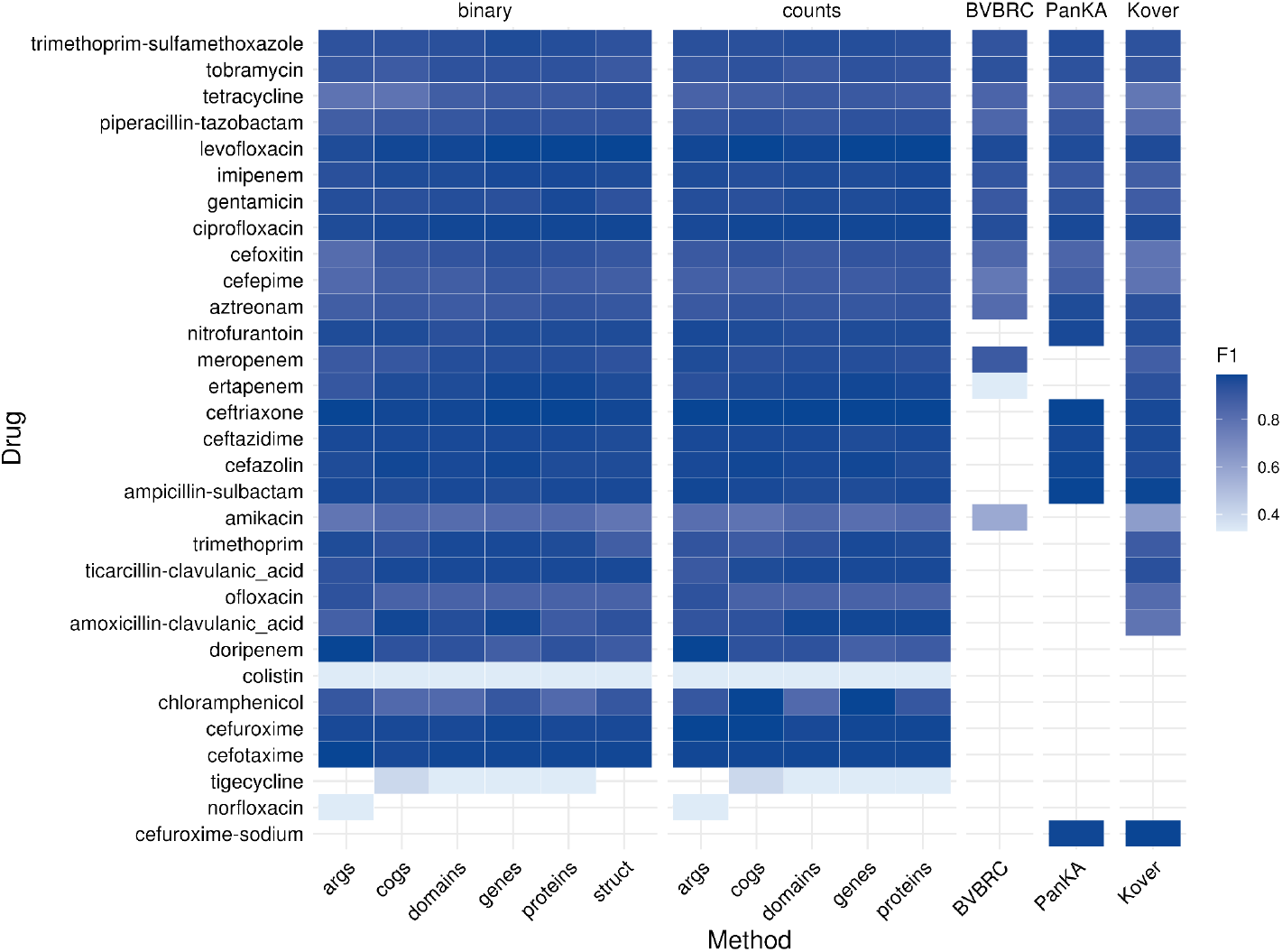
**Benchmarking the amR package suite** predictions against existing AMR phenotype prediction approaches using a selection of *Klebsiella pneumoniae* genome isolates. Heatmap colors represent F1 scores, with higher values shown in darker shades. Colors are scaled between the 5th and 95th percentiles of the F1 values for visual contrast. Antibiotics are ordered by the availability of model predictions across methods

When comparing methods, there are additional considerations beyond performance scores. First, PanKA and BV-BRC models were trained using larger datasets than what we used for our amR and Kover models, which should benefit the relative performance of the former methods. Second, while performance is an essential component of AMR prediction workflows, ease-of-use and interpretability are additional important considerations. BV-BRC as a workflow is not designed to be run by users. PanKA is designed to be run locally, but testing genomes outside their published benchmark requires manual curation of genomes, labels, and updating of their Python notebooks. Likewise, Kover is designed to identify genotype-to-phenotype associations as an ML workflow, but requires users to curate their input data and phenotypes. Its outputs include the *k*-mer rules that drive performance, offering insights into the biological features behind the models. Of these, only our amR suite offers a start-to-finish data curation, processing, ML, and visualization package, combining thorough analysis, good model performance, and an interactive dashboard to explore and understand the biology behind the model.

#### Case study: *Shigella sonnei*

To demonstrate, we applied the *amR* workflow to *Shigella sonnei*, one of the most frequent causes of dysentery.^29^ AMR is known to arise quickly in *Shigella* spp., and resistant strains can facilitate the spread of ARGs worldwide and to other bacterial pathogens.^29^ Despite its impacts and serious threat labeling by the CDC and WHO, AMR modeling in *S. sonnei* is scant. BV-BRC included data for 2,021 isolates and 43 drugs from 17 different classes. Surprisingly, the genomes included were from the 1940s to 2023 and covered around 48 countries. Ampicillin (AMP), a broad-spectrum penicillin, has the greatest number of tested genomes (∼2000) with a resistance/susceptible ratio of ∼0.5 (**Figure 3A; Table S2**). The ML models trained across feature scales (genes, proteins, domains, structural variants, COGs, and ARGs) for these genomes returned a median MCC > 0.8, reflecting strong performance (**Figure 3B; Table S2**). The top features contributing to the performance of these AMP models mapped back to Class A beta-lactamase enzymes, which are known to destroy the penicillin rings.^30,31^ When stratifying data into temporal and geographic sets, AMP models performed on par with whole models when trained and tested on the same sets (e.g., trained and tested only on US data), but when cross-testing from one time period to another, performance was almost random (MCC ≈ 0), indicating significant changes in AMP resistance mechanisms over time. This reflects the known evolutionary trajectory of beta-lactam AMR, as new beta-lactam antibiotic classes have been developed and novel resistance mechanisms have subsequently emerged across time.^32^

**Figure 3.**
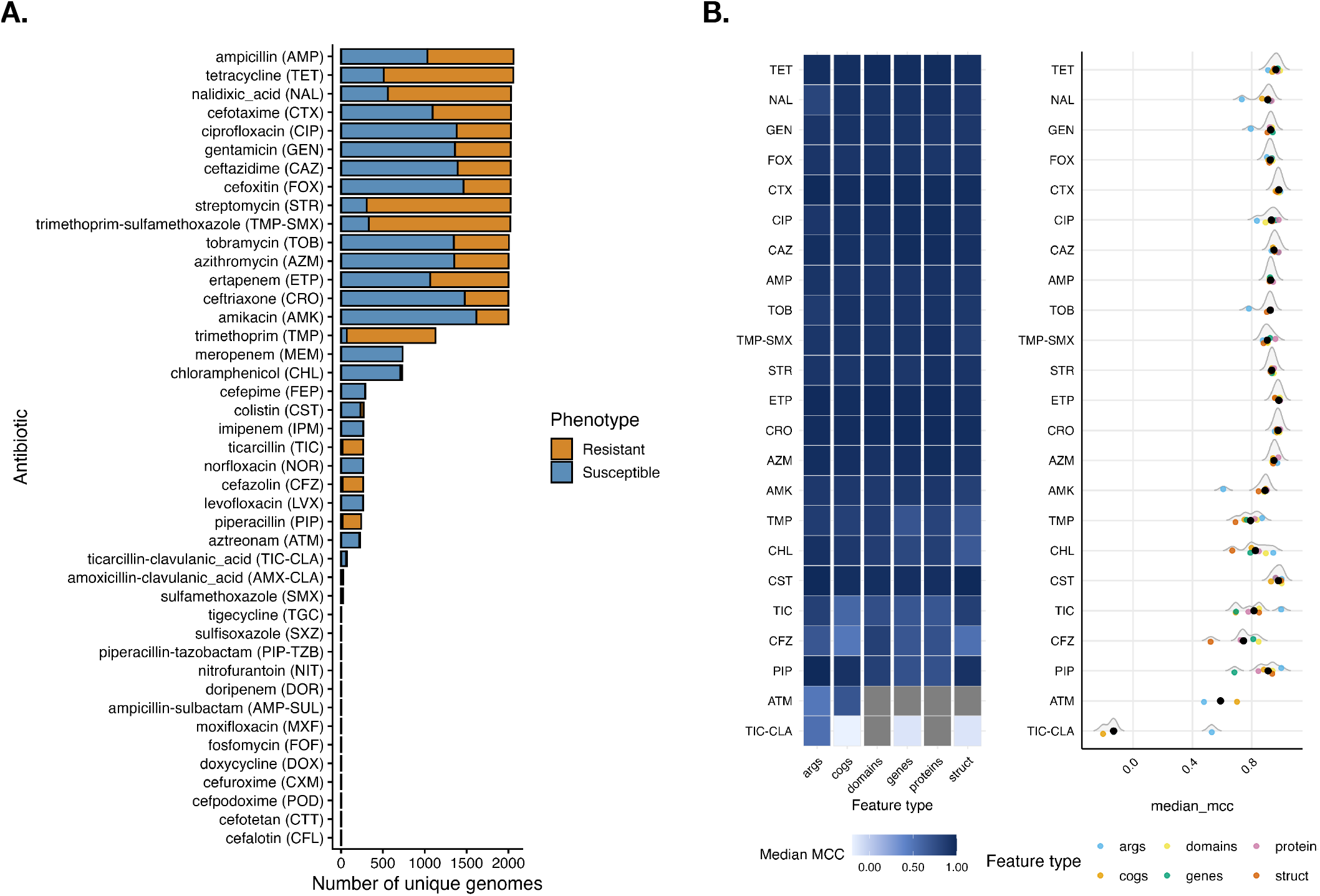
Case study: Predicting AMR in *Shigella sonnei*. **A**. The distribution of resistance phenotypes across antibiotics in *Shigella sonnei*. The stacked bars indicate the susceptible and the resistant isolate counts. **B**. The performance across drug models in *S. sonnei* is reported as the median MCC across multiple runs with different seeds. The heatmap colors range from light blue for the lowest performance to dark blue, indicating the highest performance. The accompanying ridge plot shows the distribution of median MCCs across all models for each antibiotic with points coloured by feature type

Beyond AMP, models retained high predictability across the antibiotic panel: from 23 drugs modeled, we achieved an overall median MCC of 0.89, with only 3 drugs returning median MCC values below 0.7. The best-predicted agent was cefotaxime (MCC=0.98), and at the class level, carbapenems (MCC=0.99) (**Figure 3B**). Only two antibiotics failed: aztreonam, a monobactam (MCC=-0.03), and a β-lactamase-inhibitor combination, ticarcillin-clavulanic_acid (MCC=-0.13). Notably, the poor performance reflects the skewed phenotype balance with predominance of susceptible isolates (**Figure 3A**). The top predictive features suggest well-predicted drugs were driven by recognized resistance genes (e.g., *bla* for cephalosporins, *tetA* for tetracycline), whereas the failing aztreonam models drew only on uncharacterized accessory-genome clusters. Across molecular scales, protein- and domain-level features performed best, while curated AMR genes (ARGs) alone were the weakest representation. From this, we can determine that including the broader context of other genomic feature scales improves predictions beyond known resistance genes alone.

## Discussion and Conclusion

The *amR* package suite provides a comprehensive, user-friendly, programmatic framework to predict AMR, with documented success for the notorious ESKAPE pathogens.^33^ *amR* empowers researchers to tackle biological questions that were previously inaccessible without custom workflows and computational expertise. By simplifying the setup with easy-to-use functions in R and Dockerized data and software dependencies, *amR* opens ML modeling of bacterial AMR to a wide audience. Users with basic R experience on consumer hardware can now design their own ML modeling experiments, complete data curation, processing, modeling, and visualization for a species of interest in a workday, and answer their own specific biological questions. Which poorly characterized genes drive unexplained resistance phenotypes in specific species-drug combinations? Many resistance phenotypes remain only partially explained by known AMR determinants, particularly in less-studied species and drug combinations; hence, systematic comparison of genomic features and resistance outcomes can help identify candidate genes and genomic regions associated with these unexplained patterns. How do geographic isolation sources or temporal collection periods shape resistance signatures? While it is understood that AMR patterns vary geographically and temporally,^1,34,35^ these differences are rarely modeled. By allowing users to stratify and model across time and space, we enable easier exploration of these important frontiers for emerging AMR mechanisms and those that span countries and species. What conserved protein domains or gene clusters predict AMR or convergence towards MDR across bacterial species? Similar AMR phenotypes often arise across distantly related bacterial taxa despite differences in genomic background. Examining shared protein domains and co-occurring gene clusters across species through species-agnostic modeling may reveal conserved functional mechanisms underlying MDR convergence.

Despite the BV-BRC data dependency and the large datasets associated with some bacterial species, *amR*’s modular design ensures broad applicability. Planned enhancements, including additional ML algorithms, transfer learning, and expansion to NCBI as a data source, will expand the *amR* suite’s discovery potential further.

Available at https://github.com/JRaviLab/amR with comprehensive vignettes, the *amR* suite transforms AMR ML research from black-box prediction to an out-of-the-box hypothesis generation workflow, better equipping the global research community to anticipate and counter resistance evolution. While our package suite democratizes AMR modeling, our codebase could be easily adapted to any microbial genotype-to-phenotype prediction problem.

## Acknowledgments

We would like to thank Katerina Terwelp for reviewing our manuscript and its accompanying dashboard during development. Our project relied on the numerous resources shared through BV-BRC’s database and command-line interface tools. This work utilized the Alpine high-performance computing resource at the University of Colorado Boulder. Alpine is jointly funded by the University of Colorado Boulder, the University of Colorado Anschutz, Colorado State University, and the National Science Foundation (award 2201538).

## Funding

All authors were funded by the NIH NIAID grant 1U01AI176414 and start-up funds from CU Anschutz awarded to JR. EPB was partially funded by the NIH NLM fellowship T15LM009451.

## Author contributions

Conceptualization: JR; Methodology: EPB, AG, EAB, CKV, EPW, JR; Software: EPB, AG, EAB, APM, CKV, EPW, RLL, DAM, JR; Validation: EPB, AG, CKV; Formal analysis: EPB, AG, EAB, EPW; Investigation: EPB, AG, EPW, CKV, APM; Resources: JR; Data curation: AG, EPB; Writing original draft: EPB, AG, CKV, EAB, JR; Writing review & editing: AG, EPB, CKV, EAB, APM, JR; Visualization: AG, EAB, EPB, CKV, RLL, JR; Supervision: JR; Project administration: JR; Funding acquisition: JR, EPB.

## Data availability

The amR package is GitHub-installable (through `devtools/remotes`) via the main package suite: https://github.com/jravilab/amR. The package has been submitted to Bioconductor for community review in parallel, ensuring long-term maintenance and discoverability. The individual packages can be installed from: https://github.com/jravilab/amRdata, https://github.com/jravilab/amRml, https://github.com/jravilab/amRviz. All packages are released under the BSD 3-Clause license.

## Conflict of interest

We have no conflict of interest to declare.

## Notes

https://github.com/JRaviLab/amR

### Competing Interest Statement

The authors have declared no competing interest.

## Bibliography

1. GBD 2021 Antimicrobial Resistance Collaborators. Global burden of bacterial antimicrobial resistance 1990-2021: a systematic analysis with forecasts to 2050. Lancet Lond. Engl. 404, 1199–1226 (2024).

2. de la Lastra, J. M. P., Wardell, S. J. T., Pal, T., de la Fuente-Nunez, C. & Pletzer, D. From Data to Decisions: Leveraging Artificial Intelligence and Machine Learning in Combating Antimicrobial Resistance – a Comprehensive Review. J. Med. Syst. 48, 71 (2024).

3. Anahtar, M. N., Yang, J. H. & Kanjilal, S. Applications of Machine Learning to the Problem of Antimicrobial Resistance: an Emerging Model for Translational Research. J. Clin. Microbiol. 59, 10.1128/jcm.01260-20 (2021).

4. Ghosh, A., Vang, C. K., Brenner, E. P. & Ravi, J. Unlocking antimicrobial resistance with multiomics and machine learning. Trends Microbiol. 33, 1048–1051 (2025).

5. Ismail, S. M. & Fayed, S. H. Antimicrobial Resistance Prediction in Salmonella enterica Using Frequency Chaos Game Representation and ResNet-18. Preprint at 10.1101/2025.09.13.675356 (2025).

6. Tai, H. Cross-Species Antimicrobial Resistance Prediction from Genomic Foundation Models. Preprint at 10.48550/ARXIV.2603.11141 (2026).

7. Hyun, J. C., Monk, J. M., Szubin, R., Hefner, Y. & Palsson, B. O. Global pathogenomic analysis identifies known and candidate genetic antimicrobial resistance determinants in twelve species. Nat. Commun. 14, 7690 (2023).

8. Drouin, A., Letarte, G., Raymond, F., Marchand, M., Corbeil, J. & Laviolette, F. Interpretable genotype-to-phenotype classifiers with performance guarantees. Sci. Rep. 9, 4071 (2019).

9. Aytan-Aktug, D., Clausen, P. T. L. C., Bortolaia, V., Aarestrup, F. M. & Lund, O. Prediction of Acquired Antimicrobial Resistance for Multiple Bacterial Species Using Neural Networks. mSystems 5, e00774–19 (2020).

10. Aytan-Aktug, D., Nguyen, M., Clausen, P. T. L. C., Stevens, R. L., Aarestrup, F. M., Lund, O. & Davis, J. J. Predicting Antimicrobial Resistance Using Partial Genome Alignments. mSystems 6, e00185–21 (2021).

11. Do, V. H., Nguyen, V. S., Nguyen, S. H., Le, D. Q., Nguyen, T. T., Nguyen, C. H., Ho, T. H., Vo, N. S., Nguyen, T., Nguyen, H. A. & Cao, M. D. PanKA: Leveraging population pangenome to predict antibiotic resistance. iScience 27, 110623 (2024).

12. Shukla, M., Wattam, A. R., Aleman, A., Bhattacharya, R., Bowers, N., Brettin, T., Capria, A., Chia, N., Cucinell, C., Davis, J. J., Dempsey, D. M., Dickerman, A., Dietrich, E. M., Gokdemir, O., Hendrickson, R. C., Kenyon, R. W., Klahn, B., Kuscuoglu, M., Lefkowitz, E. J., Ma, H., Machi, D., Macken, C., Mann, C. M., Mao, C., Nguyen, M., Olsen, G. J., Olson, R. D., Overbeek, R., Owens, S. M., Parrello, B., Poretsky, R., Pusch, G. D., Ramanathan, A., Sheriff, J. T., Singh, I., Stewart, L., VanOeffelen, M., Vonstein, V., Warren, A. S., Wilke, A., Zmasek, C. M., Zuniga, A. & Stevens, R. L. BV-BRC: a unified bacterial and viral bioinformatics resource with expanded functionality and AI integration. Nucleic Acids Res. 54, D715–D723 (2026).

13. Alcock, B. P., Huynh, W., Chalil, R., Smith, K. W., Raphenya, A. R., Wlodarski, M. A., Edalatmand, A., Petkau, A., Syed, S. A., Tsang, K. K., Baker, S. J. C., Dave, M., McCarthy, M. C., Mukiri, K. M., Nasir, J. A., Golbon, B., Imtiaz, H., Jiang, X., Kaur, K., Kwong, M., Liang, Z. C., Niu, K. C., Shan, P., Yang, J. Y. J., Gray, K. L., Hoad, G. R., Jia, B., Bhando, T., Carfrae, L. A., Farha, M. A., French, S., Gordzevich, R., Rachwalski, K., Tu, M. M., Bordeleau, E., Dooley, D., Griffiths, E., Zubyk, H. L., Brown, E. D., Maguire, F., Beiko, R. G., Hsiao, W. W. L., Brinkman, F. S. L., Van Domselaar, G. & McArthur, A. G. CARD 2023: expanded curation, support for machine learning, and resistome prediction at the Comprehensive Antibiotic Resistance Database. Nucleic Acids Res. 51, D690–D699 (2023).

14. Florensa, A. F., Kaas, R. S., Clausen, P. T. L. C., Aytan-Aktug, D. & Aarestrup, F. M. ResFinder – an open online resource for identification of antimicrobial resistance genes in next-generation sequencing data and prediction of phenotypes from genotypes. Microb. Genomics 8, (2022).

15. Feldgarden, M., Brover, V., Fedorov, B., Haft, D. H., Prasad, A. B. & Klimke, W. Curation of the AMRFinderPlus databases: applications, functionality and impact. Microb. Genomics 8, (2022).

16. Tonkin-Hill, G., MacAlasdair, N., Ruis, C., Weimann, A., Horesh, G., Lees, J. A., Gladstone, R. A., Lo, S., Beaudoin, C., Floto, R. A., Frost, S. D. W., Corander, J., Bentley, S. D. & Parkhill, J. Producing polished prokaryotic pangenomes with the Panaroo pipeline. Genome Biol. 21, 180 (2020).

17. Fu, L., Niu, B., Zhu, Z., Wu, S. & Li, W. CD-HIT: accelerated for clustering the next-generation sequencing data. Bioinforma. Oxf. Engl. 28, 3150–3152 (2012).

18. Paysan-Lafosse, T., Andreeva, A., Blum, M., Chuguransky, S. R., Grego, T., Pinto, B. L., Salazar, G. A., Bileschi, M. L., Llinares-López, F., Meng-Papaxanthos, L., Colwell, L. J., Grishin, N. V., Schaeffer, R. D., Clementel, D., Tosatto, S. C. E., Sonnhammer, E., Wood, V. & Bateman, A. The Pfam protein families database: embracing AI/ML. Nucleic Acids Res. 53, D523–D534 (2025).

19. Dibrova, D. V., Konovalov, K. A., Perekhvatov, V. V., Skulachev, K. V. & Mulkidjanian, A. Y. COGcollator: a web server for analysis of distant relationships between homologous protein families. Biol. Direct 12, 29 (2017).

20. Galperin, M. Y., Vera Alvarez, R., Karamycheva, S., Makarova, K. S., Wolf, Y. I., Landsman, D. & Koonin, E. V. COG database update 2024. Nucleic Acids Res. 53, D356–D363 (2025).

21. Chang, W., Cheng, J., Allaire, J. J., Xie, Y. & McPherson, J. Shiny: Web Application Framework for R. (2019).

22. Olson, R. D., Assaf, R., Brettin, T., Conrad, N., Cucinell, C., Davis, J. J., Dempsey, D. M., Dickerman, A., Dietrich, E. M., Kenyon, R. W., Kuscuoglu, M., Lefkowitz, E. J., Lu, J., Machi, D., Macken, C., Mao, C., Niewiadomska, A., Nguyen, M., Olsen, G. J., Overbeek, J. C., Parrello, B., Parrello, V., Porter, J. S., Pusch, G. D., Shukla, M., Singh, I., Stewart, L., Tan, G., Thomas, C., VanOeffelen, M., Vonstein, V., Wallace, Z. S., Warren, A. S., Wattam, A. R., Xia, F., Yoo, H., Zhang, Y., Zmasek, C. M., Scheuermann, R. H. & Stevens, R. L. Introducing the Bacterial and Viral Bioinformatics Resource Center (BV-BRC): a resource combining PATRIC, IRD and ViPR. Nucleic Acids Res. 51, D678–D689 (2023).

23. Sean R. E. HMMER: biosequence analysis using profile hidden Markov models. (2023).

24. Mistry, J., Chuguransky, S., Williams, L., Qureshi, M., Salazar, G. A., Sonnhammer, E. L. L., Tosatto, S. C. E., Paladin, L., Raj, S., Richardson, L. J., Finn, R. D. & Bateman, A. Pfam: The protein families database in 2021. Nucleic Acids Res. 49, D412–D419 (2021).

25. Jones, P., Binns, D., Chang, H.-Y., Fraser, M., Li, W., McAnulla, C., McWilliam, H., Maslen, J., Mitchell, A., Nuka, G., Pesseat, S., Quinn, A. F., Sangrador-Vegas, A., Scheremetjew, M., Yong, S.-Y., Lopez, R. & Hunter, S. InterProScan 5: genome-scale protein function classification. Bioinforma. Oxf. Engl. 30, 1236–1240 (2014).

26. Raasveldt, M. & Mühleisen, H. DuckDB: an Embeddable Analytical Database. in Proceedings of the 2019 International Conference on Management of Data 1981–1984 (Association for Computing Machinery, New York, NY, USA, 2019). doi:10.1145/3299869.3320212.

27. Merkel, D. Docker: lightweight linux containers for consistent development and deployment. Linux J. 2014, 2 (2014).

28. Kuhn, M. & Wickham, H. Tidymodels: a collection of packages for modeling and machine learning using tidyverse principles. (2020).

29. Kotloff, K. L., Riddle, M. S., Platts-Mills, J. A., Pavlinac, P. & Zaidi A. K. M. Shigellosis. The Lancet 391, 801–812 (2018).

30. Bush, K. Past and Present Perspectives on β-Lactamases. Antimicrob. Agents Chemother. 62, e01076–18 (2018).

31. He, Y., Lei, J., Pan, X., Huang, X. & Zhao, Y. The hydrolytic water molecule of Class A β-lactamase relies on the acyl-enzyme intermediate ES* for proper coordination and catalysis. Sci. Rep. 10, 10205 (2020).

32. Fröhlich, C., Chen, J. Z., Gholipour, S., Erdogan, A. N. & Tokuriki, N. Evolution of β-lactamases and enzyme promiscuity. Protein Eng. Des. Sel. 34, gzab013 (2021).

33. Ghosh, A., Brenner, E. P., Vang, C. K., Wolfe, E. P., Boyer, E. A., Lesiyon, R. L., Manpearl, K. R., Sridhar, V., Burke, J. T., Krol, J. D., Bilodeaux, J. M. & Ravi, J. From sequence to signature: Machine learning uncovers multiscale feature landscapes that predict AMR across ESKAPE pathogens. Preprint at 10.1101/2025.07.03.663053 (2025).

34. Keely, S. P., Brinkman, N. E., Wheaton, E. A., Jahne, M. A., Siefring, S. D., Varma, M., Hill, R. A., Leibowitz, S. G., Martin, R. W., Garland, J. L. & Haugland, R. A. Geospatial Patterns of Antimicrobial Resistance Genes in the US EPA National Rivers and Streams Assessment Survey. Environ. Sci. Technol. 56, 14960–14971 (2022).

35. Legenza, L., McNair, K., Gao, S., Lacy, J. P., Olson, B. J., Fritsche, T. R., Schulz, L. T., LaMuro, S., Spray-Larson, F., Siddiqui, T. & Rose, W. E. A geospatial approach to identify patterns of antibiotic susceptibility at a neighborhood level in Wisconsin, United States. Sci. Rep. 13, 7122 (2023).

